# Monovalent Ion Effect on Liquid–Liquid Phase Separation of Aqueous Polyphosphate–Salt Mixtures

**DOI:** 10.1101/2024.05.01.592046

**Authors:** Tomohiro Furuki, Azusa Togo, Hatsuho Usuda, Tomohiro Nobeyama, Atsushi Hirano, Kentaro Shiraki

## Abstract

Polyphosphate (polyP) is one of the most conserved biomacromolecules and can form aggregates, such as polyP granules in bacteria, which are generated through liquid–liquid phase separation (LLPS). Studies have examined the mechanism of polyP aggregation using LLPS systems containing artificial polyP molecules as aggregation system models, where LLPS is typically induced by multivalent salts and polyelectrolytes. Although the typical concentrations of monovalent ions in living cells are approximately 100 times higher than those of divalent ions, the effects of monovalent ions on the LLPS of polyP solutions are little known. This study demonstrated that submolar NaCl induces LLPS of polyP solutions, whereas other monovalent salts did not at the same concentrations. Small-angle X-ray scattering measurements revealed that NaCl significantly stabilizes the intermolecular association of polyP, inducing LLPS. These findings suggest that the modulation of monovalent ion concentrations is an underlying mechanism of polyP aggregate formation/deformation within living cells.

**TOC GRAPHIC:** 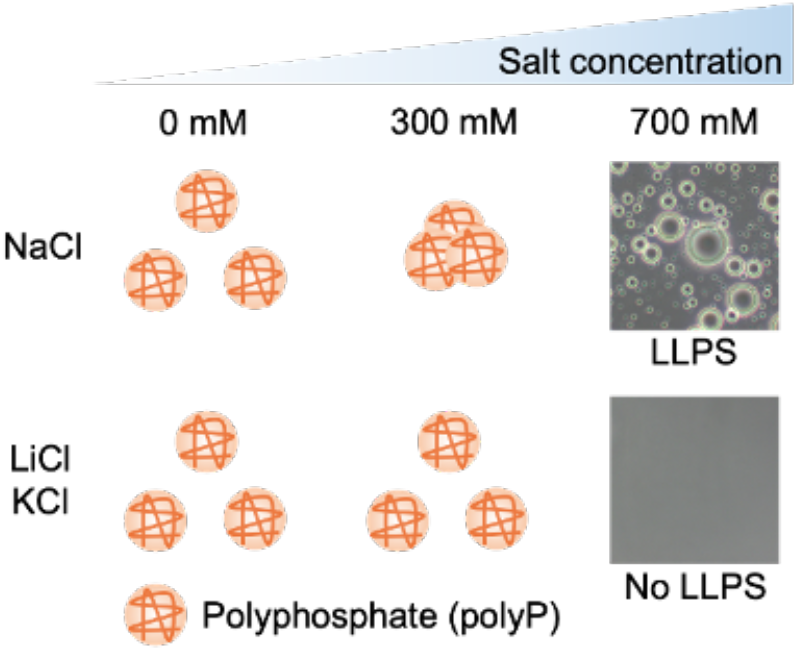

---

Polyphosphate (polyP) is one of the most conserved biomacromolecules.^1^ It has a linear structure, typically consisting of 3–1000 phosphate units. The presence of polyP in living cells was first identified as a volutin granule in the early 20th century.^2^ Volutin granules are an aggregated form of polyP. Since then, polyP aggregates have been found in living cells, such as acidocalcisomes in Trypanosomatida^3^ and platelet-dense granules in humans.^4^ PolyP granules are one of the most investigated aggregates and are formed in many bacterial cells through liquid–liquid phase separation (LLPS) under stress such as nutrient starvation.^5^ However, the physicochemical mechanism of polyP granule formation remains unclear.

The mechanism of polyP granule formation has been examined using polyP solutions containing multivalent cations and polyelectrolytes as in vitro LLPS models because polyP granules are abundant in Mg^2+^ ^6, 7^. In fact, MgCl_2_ and KCl at the same concentrations as those in *Escherichia coli* induce LLPS of polyP solutions.^8^ Mn^2+^ and arginine–glutamic acid–arginine peptides induce LLPS of polyP solutions, which confer radiotolerance to coexisting proteins.^9^ Positively charged +36GFP induces LLPS of polyP solutions.^10^ In addition to multivalent cations and polyelectrolytes, monovalent cations are also present in cells, typically at concentrations of several hundred millimolars. However, their effects on LLPS of polyP solutions are not well known.

This study demonstrated that submolar NaCl induces LLPS of polyP solutions, whereas other alkali metal salts, namely, KCl, RbCl, and CsCl, did not at the same concentrations. Small-angle X-ray scattering (SAXS) measurements demonstrated that NaCl stabilizes the intermolecular association of polyP, which can contribute to its LLPS-inducing effect. These findings suggest that monovalent cations regulate polyP aggregation in living cells.

Submolar NaCl induced LLPS of polyP solutions, where LLPS was defined as the generation of spherical-shaped droplets observed under a microscope. Figure 1a shows a typical droplet containing polyP with 0.9 M NaCl; notably, polyP was stained by 4’,6-diamidino-2-phenylindole (DAPI), a yellow fluorescence compound used to detect polyP with ≥15 phosphate units^11^; a DAPI–polyP complex exhibits fluorescence with a peak wavelength of approximately 525 nm (Figure S1).^12, 13^ PolyP droplets were fused (Figure 1b), indicating that they have a liquid-like property. Figure 1c shows the phase diagram of LLPS for polyP with 60–70 phosphate units, referred to as polyP_60_, at different NaCl concentrations. LLPS of polyP_60_ at a phosphate unit molarity of 1 mM (referred to as 1 mM-P) was induced at ≥1 M NaCl. LLPS of polyP_60_ solutions at 10 and 100 mM-P was induced at ≥0.7 M NaCl. In polyP with ≥700 phosphate units (polyP_700_), LLPS was induced even at lower NaCl concentrations, i.e., 0.5 M (Figure 1d). In contrast, in polyP with 10–15 phosphate units (polyP_10_), LLPS was not induced at 0–3.4 M NaCl (Figure S2). The reduced critical concentrations of NaCl in polyP_700_ can be accounted for by the Flory–Huggins theory, which indicates that a higher degree of polymerization lowers the critical concentration for LLPS.^14^ In conclusion, submolar NaCl can induce LLPS of polyP solutions, which depends on polyP’s molecular length and concentration.

**Figure 1.**
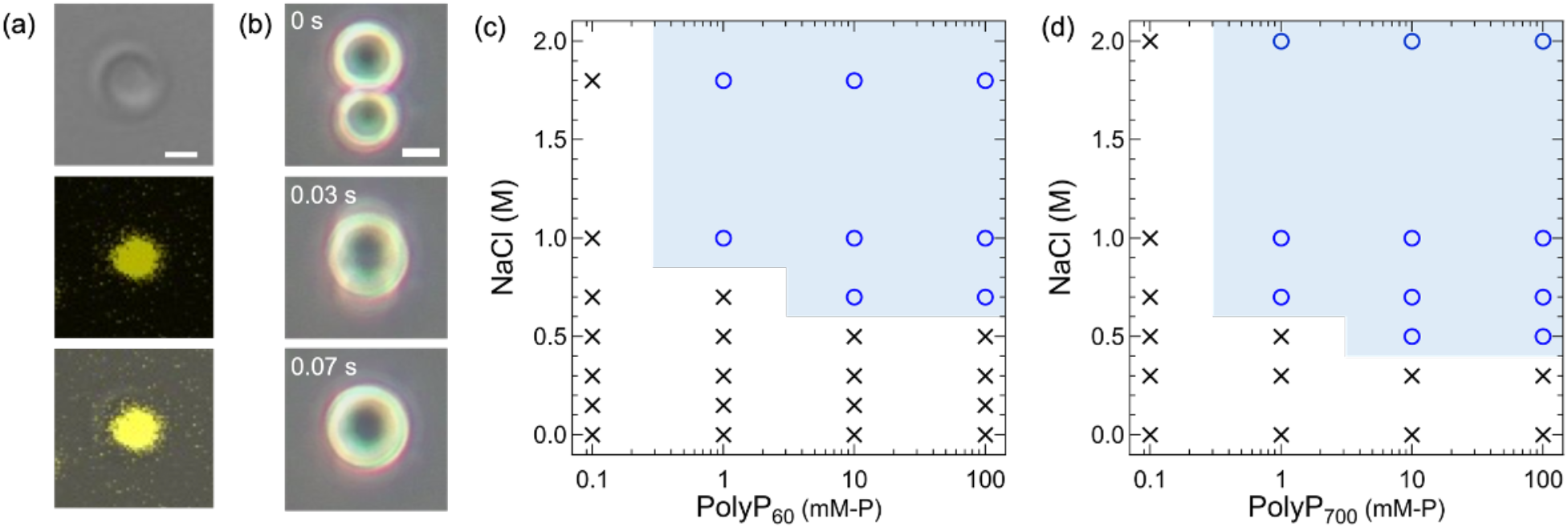
(a) Fluorescence image of a polyP_60_ droplet formed through LLPS at a concentration of 12.5 mM-P with 0.9 M NaCl. From top to bottom: transmission image, fluorescence image of the DAPI– polyP complex (yellow), and merged image (scale bar, 1 μm). (b) Snapshot of the fusion event of polyP_60_ droplets (scale bar, 5 μm). (c, d) Phase diagram of polyP_60_ (c) and polyP_700_ (d) solutions with NaCl. Circles and crosses represent LLPS occurrence and no LLPS occurrence, respectively. The unit mM-P corresponds to phosphate unit molarity in millimolars.

Other alkali metal salts, namely, LiCl, KCl, RbCl, and CsCl, were less effective or ineffective in contrast to NaCl. Among these salts, only LiCl induced LLPS of the 10 mM-P polyP_60_ solution at ≥4.5 M concentrations (Figure 2a); note that 9 M LiCl induced the formation of polyP aggregates with non-spherical shapes (Figure S3).

**Figure 2.**
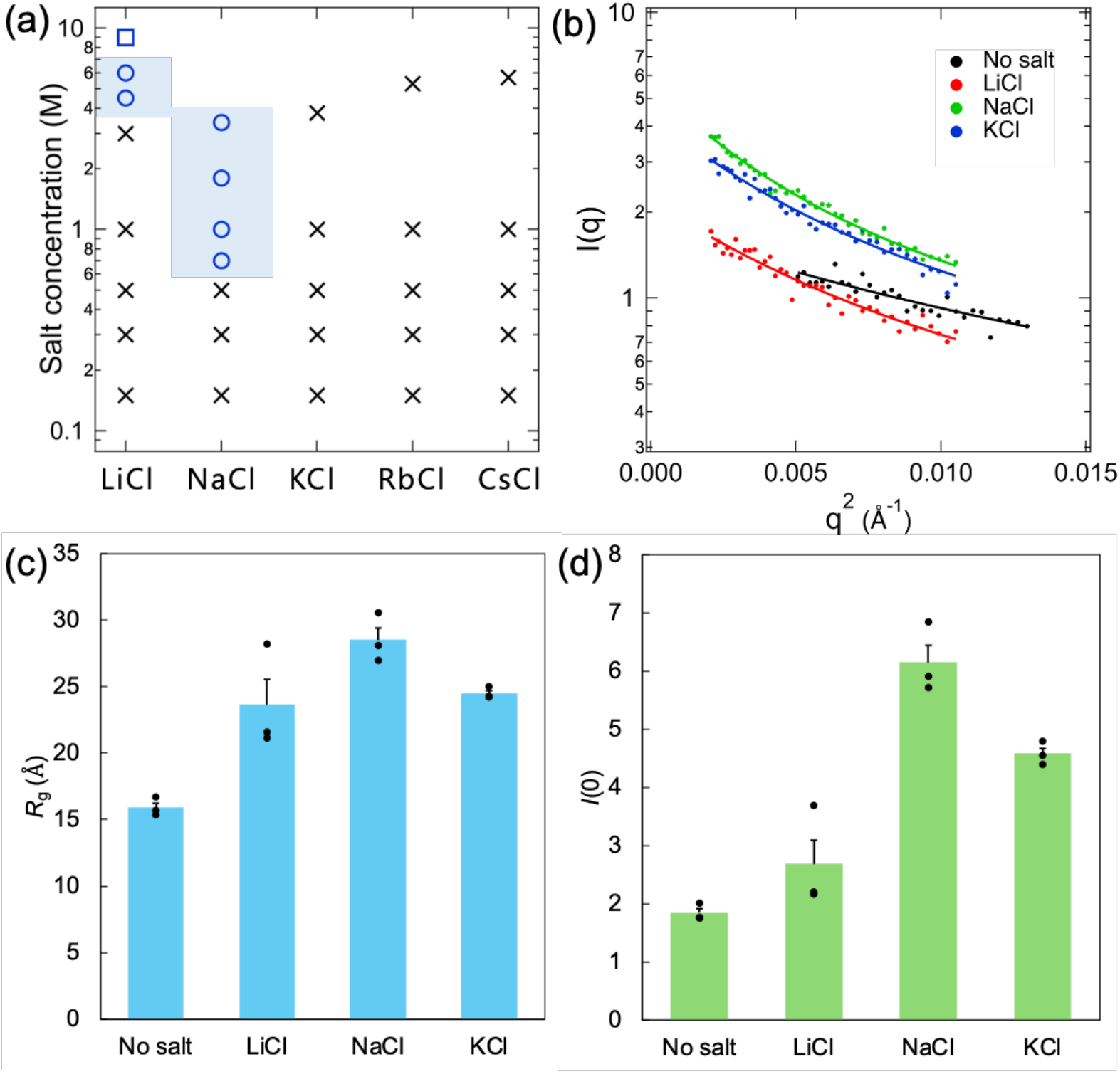
(a) Phase diagram of 10 mM-P polyP_60_ solutions with monovalent salts. Circles, crosses, and squares represent LLPS occurrence, no LLPS occurrence, and aggregate formation, respectively. (b) Fit of the Debye function to the SAXS profiles of 100 mM-P polyP_60_ solutions containing 0 and 300 mM monovalent salts. (c, d) Values of *R*_g_ (c) and *I*(0) (d) obtained by fitting. Triplicate data are plotted on the bar. Error bars represent standard errors for the triplicate data.

In general, LLPS is induced by intermolecular interactions. Thus, the significant effect of NaCl on the induction of LLPS of polyP is assumed to be ascribed to the microscopic association of polyP molecules. This assumption was verified by SAXS for polyP_60_ with and without 300 mM monovalent salts, i.e., LiCl, NaCl, and KCl. SAXS data of scattered intensity *I* as a function of the scattering vector *q* was fitted to the following Debye function^15, 16^ in the region of 0 < *qR*_g_ < 3 (Figure 2b):

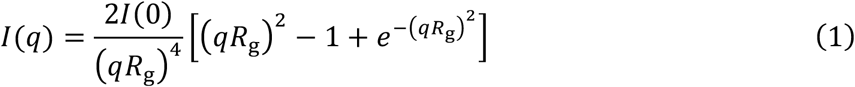

where *R*_g_ is the radius of gyration of polyP_60_, and *I*(0) is a parameter,

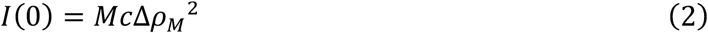

where *M* is the single-particle mass, *c* presents the weight concentration of the solute, and Δ*ρ*_*M*_ indicates the scattering contrast per mass, which is proportional to the number of electrons.^17^ This function has been widely used for unstructured biomacromolecules, such as intrinsically disordered proteins and single-stranded DNA.^15, 16, 18^ PolyP_60_ showed the largest *R*_g_ in the presence of NaCl (Figure 2c). This result matches that of the phase diagram shown in Figure 2a. The significantly high *I*(0) value with NaCl indicates an increase in single-particle mass (Figure 2d), suggesting an increase in the apparent molecular weight of polyP_60_ through intermolecular association. In contrast, the difference in the *I*(0) value between KCl and LiCl can be simply accounted for by the larger number of electrons in K^+^ ion than in Li^+^ ion. These results indicate that NaCl effectively stabilizes the intermolecular association of polyP, leading to LLPS.

The LLPS-inducing effects of Li^+^ and Na^+^ ions can be attributed to larger surface charge densities than those for the other cations, which can cause stronger electrostatic interactions with negatively charged polyP. Although Li^+^ ions have a smaller ion radius and larger surface charge density than Na^+^ ions, the effective surface charge densities of Li^+^ ions in aqueous solutions are lower than those of Na^+^ ions, which is attributed to the formation of tightly bound first hydration shells.^19^ Thus, the lowest critical NaCl concentration among the five alkali metal salts is ascribable to the strongest electrostatic interactions with polyP. Incidentally, LLPS was dependent on the nature of anions (Figures S2–S4). NaSCN and NaI induced LLPS of the polyP_60_ solution at lower concentrations than NaCl (Figure S4); note that 7.5 M NaSCN induced the formation of polyP aggregates with nonspherical shapes (Figure S3). The order of the critical concentrations for LLPS among anions matched with the Hofmeister series Cl^−^, I^−^, and SCN^−^.^20, 21^

The potential contributions of hydrophobic interactions in LLPS were examined by determining the occurrence of LLPS with and without 1,6-hexanediol, which is known for reducing hydrophobic interactions^22^. LLPS of a 10 mM-P polyP_60_ solution was induced with and without 10% w/v 1,6-hexanediol in the presence of 1 M NaCl (Figure S5), suggesting that hydrophobic interaction does not dominantly contribute to the LLPS of the polyP_60_ solution. In general, LLPS is more readily induced at higher temperatures for hydrophobic interaction-induced LLPS, and such a hydrophobic interaction is stronger at higher temperatures.^23^ In experiments with varying temperatures, 0.47 M NaCl induced LLPS of 100 mM-P polyP_60_ solution at 4°C but not at 25°C, which also likely supports the insignificance of hydrophobic interactions (Figure S6).

In living cells, cytosols are not a dilute system of biomolecules but highly concentrated systems that contain approximately 0.3–0.4 g mL^−1^ biomacromolecules, such as proteins and nucleic acids.^24^ To mimic such a crowded environment in cells, macromolecules called “crowders,” such as polyethylene glycol (PEG), dextran, and bovine serum albumin (BSA), have been used in the studies on LLPS of biomacromolecules.^25^ Generally, crowders promote LLPS through several mechanisms, including the volume exclusion effect, specific associative or segregative interaction with polymers, and gain of dehydration entropy.^25, 26^ Essentially, 5%–20% PEG decreases the critical concentrations of various protein solutions for LLPS.^27^ In this study, PEG with a molecular weight of approximately 6,000 and BSA decreased the critical concentrations for LLPS of the polyP solution (Figure 3). With 5% w/v PEG, LLPS of the 1–10 mM-P polyP_60_ solutions was induced at NaCl concentrations of ≥0.5 M and that of the 100 mM-P polyP_60_ solution at NaCl concentrations of ≥0.3 M (Figure 3a). With 10% w/v PEG, LLPS of polyP_60_ solutions was induced even at lower NaCl concentrations (Figure 3b). Thus,

**Figure 3.**
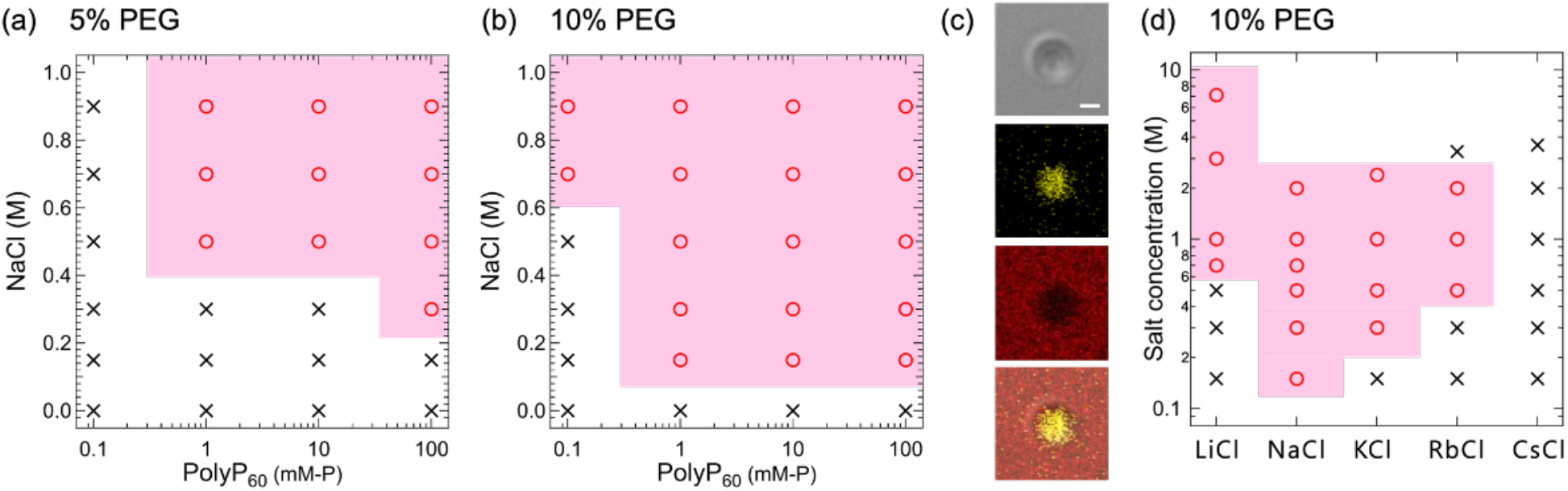
(a, b) Phase diagram of polyP_60_ solutions with NaCl and PEG at 5% w/v (a) and 10% w/v (b). (c) Fluorescent images of a polyP_60_ droplet formed through LLPS at 12.5 mM-P with of 0.9 M NaCl and 6.4% w/v PEG. From top to bottom: transmission image, fluorescent image of the DAPI– polyP complex (yellow), rhodamine–PEG (red), and merged image (scale bar, 1 μm). (d) Phase diagram of the 10 mM-P polyP_60_ solutions with monovalent salts and 10% w/v PEG. Circles and crosses represent LLPS occurrence and no LLPS occurrence, respectively. The unit mM-P corresponds to phosphate unit molarity in millimolars.

PEG can reduce the critical NaCl concentrations for LLPS to the physiological salt concentration (0.15 M). A similar effect was observed for BSA, although weaker (Figure S7). The fluorescent image of the DAPI–polyP and rhodamine–PEG complex indicated that PEG was excluded from the polyP droplet (Figure 3c), which suggests that the effect of PEG on the LLPS is attributed to the volume exclusion effect. Absorbance measurements of macroscopically separated phases confirmed the exclusion of PEG from the polyP droplet (data not shown).

PEG also enhanced the LLPS-inducing effects of other alkali metal salts. The critical concentrations of LiCl, NaCl, KCl, and RbCl for LLPS of polyP_60_ were significantly lower with 10% w/v PEG than without it (Figure 2a and 3d). This result indicates that crowders enhanced the LLPS-inducing effect of alkali metal salts. However, CsCl did not induce LLPS at 0–3.6 M concentrations even with 10% w/v PEG (Figure 3d), indicating a substantially weaker the LLPS-inducing effect of CsCl. LLPS was not induced by 3.3 M RbCl. This result can be attributed to the loss of charge balance caused by the excess amount of cation.^28^

This study mainly focused on the effect of monovalent ions on LLPS of the polyP solution. However, not only submolar monovalent ions but also millimolar divalent ions coexist in living cells.^29^ LLPS occurrence in polyP solution was further examined with 1–1500 mM MgCl_2_ (Figure S8a) and coexistence of 0–2 M NaCl and 0–20 mM MgCl_2_ (Figure S8b). Under certain conditions, NaCl and MgCl_2_ coexistence can inhibit LLPS of the polyP solution (Figure S8b). Moreover, 10% w/v PEG and 20% w/v BSA lowered the critical concentration of NaCl for the LLPS but did not lower that of MgCl_2_ (Figure S8c, d). These complicated phenomena present information about the cooperative role of monovalent and divalent metal ions in LLPS, which should be addressed in future research to better understand LLPS in living systems.

This study demonstrated that 0.5–2 M NaCl induces LLPS of the polyP solution. Conversely, other alkali metal salts, such as KCl, RbCl, and CsCl, did not induce LLPS of polyP solutions even at the same concentrations. SAXS measurements revealed that NaCl promotes the intermolecular association of polyP, which can confirm its significant LLPS-inducing effect. These findings suggest that monovalent cations can help regulate LLPS by controlling the nanoscale association of polyP, which may be possible in biological cells.

## EXPERIMENTAL METHODS

### Materials

PolyP_10_ (average chain length, 10–15), PolyP_60_ (average chain length, 60–70) and polyP_700_ (average chain length, 700–800) were purchased from Bioenex (Hiroshima, Japan). NaCl, CsCl, Na_2_SO_4_, NaH_2_PO_4_, NaHCO_3_, NaSCN, and MgCl_2_ were purchased from Kanto Chemical (Tokyo, Japan). KCl was obtained from Nacalai Tesque (Kyoto, Japan). LiCl, RbCl, NaI, and polyethylene glycol 6,000 (PEG) were acquired from Wako Pure Chemical (Osaka, Japan). DAPI and BSA were purchased from Sigma Aldrich (MO, USA). mPEG-Rhodamine B, 5k (rhodamine PEG) was bought from Biopharma PEG (MA, US). All stock solutions, except for NaHCO_3_, contained 10 mM HEPES with pH 7.4, and NaHCO_3_ stock solution contained 10 mM HEPES, with pH 8.4.

### PolyP droplet observation with different kinds of salts and crowders

A 0.1-mm thick silicon sheet with a hole measuring approximately 6 mm inserted between a silicon-coated slide glass (Matsunami Glass, Osaka, Japan) and a 0.13–0.17-mm thick cover glass (Matsunami Glass) was used as a chamber for droplet observation. An optical microscope (BA310PH, Shimadzu Rika Corporation, Kyoto, Japan) equipped with a 40× lens in a phase contrast mode was used for obtaining physical images of the samples. PolyP, salt, and crowder stock solutions, which contain 10 mM HEPES (pH 7.4), were separately prepared. BSA concentration was determined using a spectrophotometer (ND-1000 Instrument, NanoDrop Technologies, DE, USA). To achieve final concentrations of interests, PolyP, salt, and crowder stock solutions were mixed in microtubes by pipetting. The total volume of each sample was 20 μL. To form uniform droplets, the samples were vortexed immediately for 20 s, and 3.5-μL aliquots of the resulting suspensions were dropped into the hole. Droplets were observed 10–20 min after the dropping.

### Fluorescent observation of polyP droplets

Droplets were observed using a confocal laser scanning microscope (IX81, FV1000; Olympus Inc., Tokyo, Japan) equipped with a water immersion objective lens (UPLSAPO 60XW, Olympus Inc.). The final concentrations of DAPI with and without PEG were 8 and 58 μM, respectively. The final concentration of rhodamine–PEG was 0.05 mg mL^−1^. DAPI and rhodamine–PEG were excited at wavelengths of 405 and 559 nm and detected in the ranges of 570–620 and 575–675 nm, respectively. A DAPI–polyP complex emitted green-yellow fluorescence, which is different from that of DAPI and the DAPI–nucleic acid complex^12,13^. A fluorescence spectrum of the DAPI–polyP complex inside a polyP droplet excited with a 405-nm laser was obtained in the wavelength range of 420–612 nm with an 8-nm interval. To prevent fluorescence overlap, DAPI and rhodamine were excited separately. The fluorescence intensity was analyzed using the Fiji software (National Institute of Health, USA).

### SAXS

SAXS measurement was conducted using NANOPIX (Rigaku Co., Tokyo, Japan) at 40 kV and 30 mA with CuK_α_ radiation (*λ* = 0.15418 nm) at an ambient temperature. The sample-to-detector (HyPix-6000, Rigaku Co.) distance was 1299.78 mm, calibrated by silver behenate. Samples containing 100 mM-P polyP_60_, 12.5 mM HEPES (pH 7.4), and with or without 300 mM LiCl, NaCl, and KCl had been prepared on the previous day and stored at 4°C. Reference solutions containing 12.5 mM HEPES (pH 7.4), with or without 300 mM LiCl, NaCl, and KCl, were used to obtain background signals. Triplicate measurements were performed in a polyimide tube with an inner diameter of 1.6 mm (Furukawa Electric Co., Ltd., Tokyo, Japan). Each sample was subjected to the SAXS measurement for 5 min with a *q*-range of 0.04–0.11 Å^−1^. Analyses of two- and one-dimensional X-ray patterns (converted from the two-dimensional pattern) were performed using the 2DP software (Rigaku Co.). Scattering intensities were normalized using direct intensity values. To obtain the intensity originating from polyP, the intensities of air, 300 mM salt, and HEPES solution were subtracted from those of all samples. Fitting the Debye function to the data was performed by Igor Pro 9 (WaveMetrics, US).

## ASSOCIATED CONTENT

Supporting Information Available: Additional experimental data (docx).

### Notes

The authors declare no competing financial interest.

## ACKNOWLEDGEMENTS

We thank Y. Takenaka for supporting the SAXS experiments. This study was supported by Grants-in-Aid for Scientific Research from Japan Society for the Promotion of Science (Grant no. 22K18316 [K.S.] and 22K14573 [T.N.]).

