## Supporting Information for "Monovalent Ion Effect on Liquid–Liquid Phase Separation of Aqueous Polyphosphate–Salt Mixtures"

Supplementary Figures

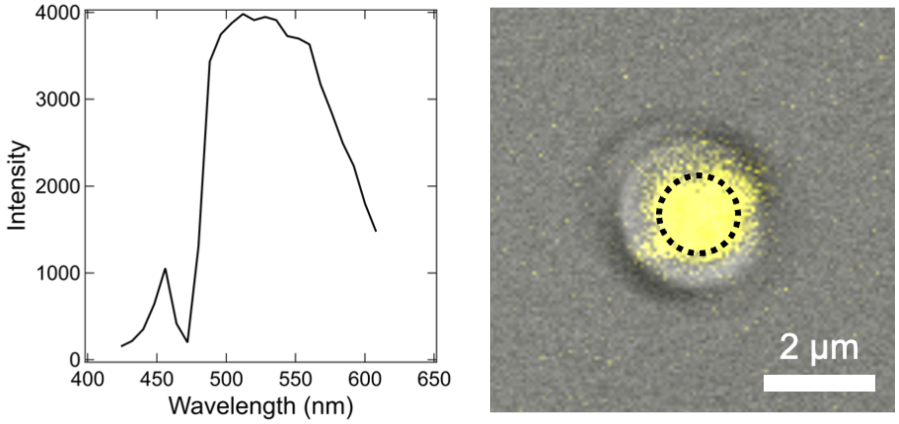


**Figure S1.** Fluorescence spectrum of the DAPI–polyP$_{60}$ complex inside a polyP$_{60}$ droplet (corresponding to the average intensity obtained within the dotted circle on the right panel) obtained using a confocal laser scanning microscope.

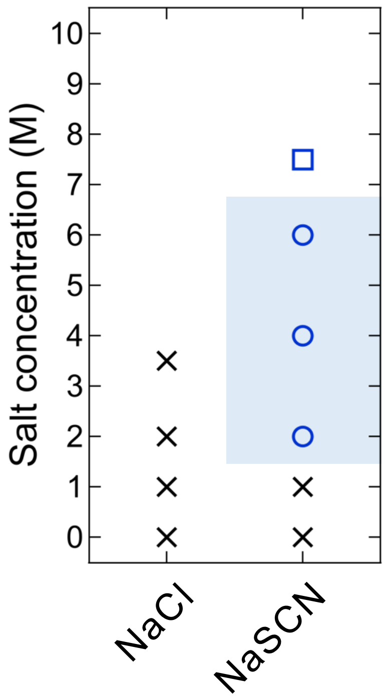

**Figure S2.** Phase diagram of the 10 mM-P polyP$_{10}$ solutions with NaCl or NaSCN. Circles, crosses, and squares represent LLPS occurrence, no LLPS occurrence, and aggregate formation, respectively.

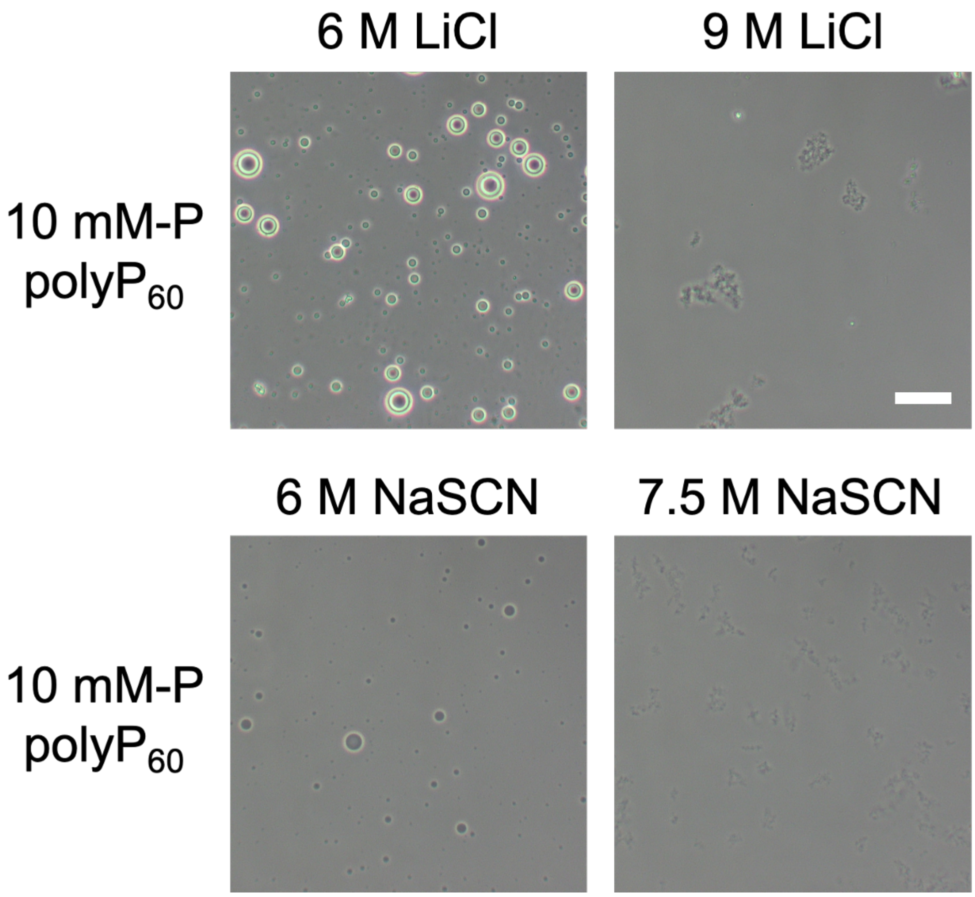


**Figure S3.** LLPS and aggregate formation of 10 mM-P polyP$_{60}$ solutions at high concentrations of LiCl and NaSCN (scale bar, 25 μm).

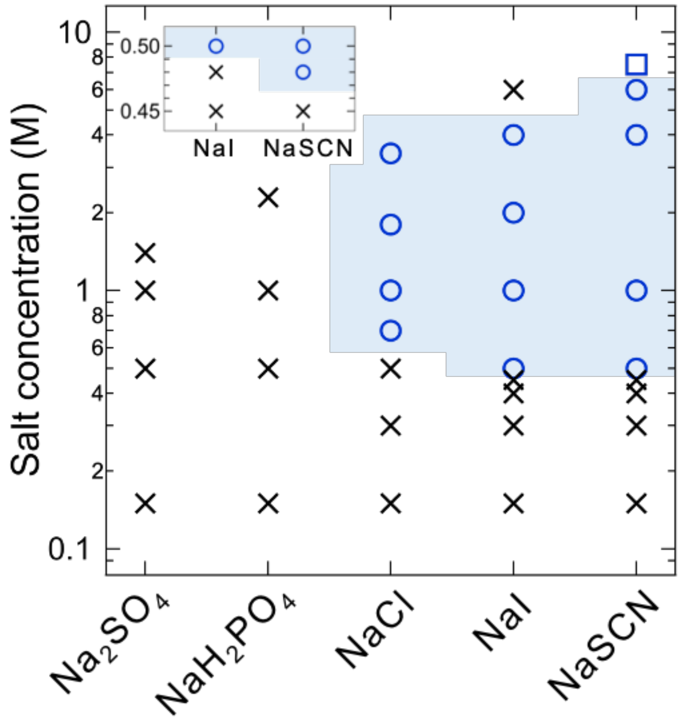


**Figure S4.** Phase diagram of 10 mM-P polyP$_{60}$ solutions with sodium salts. Circles, crosses, and squares represent LLPS occurrence, no LLPS occurrence, and aggregate formation, respectively.

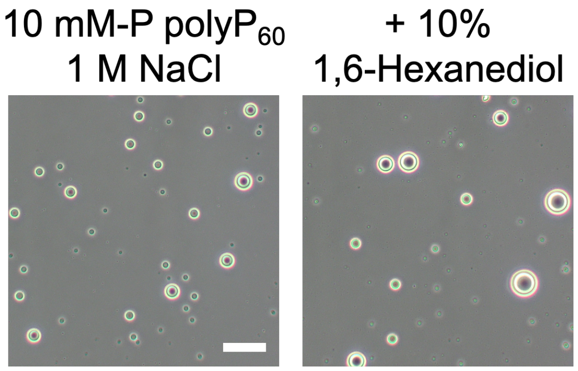


**Figure S5.** LLPS of 10 mM-P polyP$_{60}$ solutions with or without 10% w/v 1,6-hexanediol with 1 M NaCl (scale bar, 25 µm).

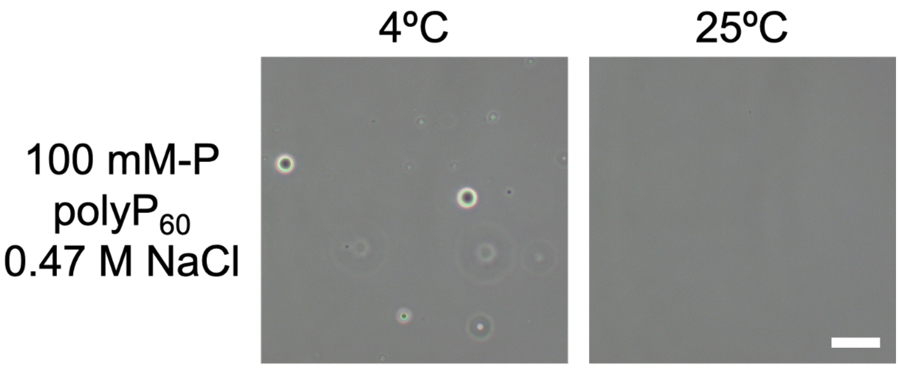


**Figure S6.** Temperature dependence of LLPS of a 100 mM-P polyP$_{60}$ solution with 0.47 M NaCl (scale bar, 25 µm).

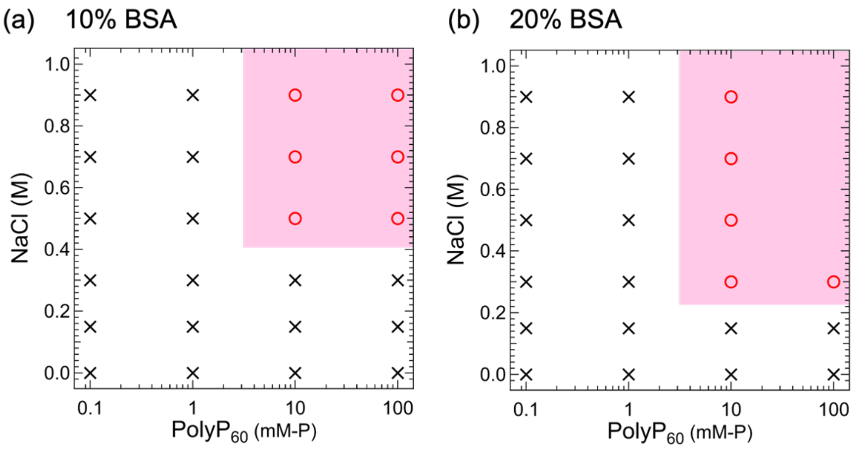


**Figure S7.** Phase diagram of the polyP$_{60}$ solutions with NaCl along with PEG at 10% w/v (a) and 20% w/v (b). Circles and crosses represent LLPS occurrence and no LLPS occurrence, respectively.

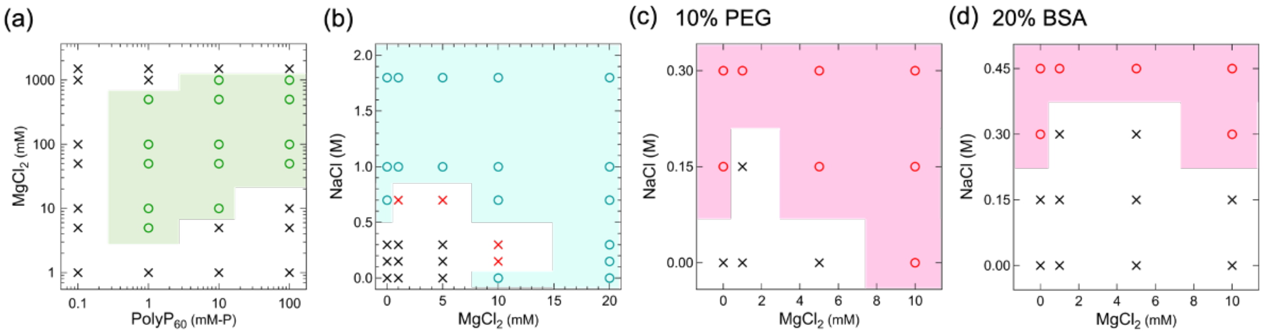


**Figure S8.** (a) Phase diagrams of polyP$_{60}$ solutions with MgCl$_2$. (b) Phase diagram of 10 mM-P polyP$_{60}$ solutions with NaCl and MgCl$_2$. Red crosses represent the conditions where LLPS was suppressed by NaCl and MgCl$_2$ coexistence. (c, d) Phase diagrams of the LLPS of 10 mM-P polyP$_{60}$ solutions with NaCl and MgCl$_2$ along with PEG at 10% w/v (c) and 20% w/v (d). Circles and crosses represent LLPS occurrence and no LLPS occurrence, respectively.
